# Neural Flip-Flops II: The Role of Cascaded Oscillators in Short-Term Memory, EEGs, and Epilepsy

**DOI:** 10.1101/2020.06.24.168419

**Authors:** Lane Yoder

## Abstract

By enabling many brain structures’ state changes, the explicit cascaded oscillators proposed here can generate the rhythmic neural activity found in EEGs. The function of such synchronization in information processing systems is timing error avoidance. The narrow requirement for the oscillator input pulse duration suggests a possible relationship to the abnormal electrical activity characteristic of epileptic seizures. Together, flip-flops and synchronization by oscillators suggest a resolution to the longstanding controversy of whether short-term memory depends on neurons firing persistently or in brief, coordinated bursts.

The proposed cascade of oscillators consists of a ring oscillator and four toggle flip-flops connected in sequence. The novel oscillator and toggle are composed of three and six neurons, respectively. Their operation depends only on minimal properties of excitatory and inhibitory inputs.

The hypothesis that cascaded oscillators produce EEG phenomena implies that the distribution of EEG frequencies is determined by just two parameters, the mean (μ_d_) and standard deviation (σ_d_) of the delay times of neurons that make up the initial oscillators in the cascades. For example, if μ_d_ and σ_d_ are measured in milliseconds, the boundary separating the alpha and beta frequency bands is

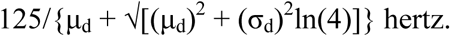

With 4 and 1.5 ms being the best available estimates for μ_d_ and σ_d_, respectively, this predicted boundary value is 14.9 Hz, which is within the range of commonly cited estimates obtained empirically from EEGs. Four and 1.5 ms also accurately predict the peaks and other boundaries of the five major EEG frequency bands.

**Significance statement:** The neuronal model proposed here implies several major aspects of electroencephalography. The matched periods of neural activity found in EEGs, and their wide distribution across the brain and across the frequency spectrum, follow from selective pressure for a biologically useful function: timing error avoidance for diverse brain functions in the trade-off between speed and accuracy. This activity can be achieved with a simple organization of synaptic connections and minimal neuron capabilities of excitation and inhibition. The multimodal distribution of EEG frequencies is an explicit function of the mean and variance of neuron delay times. The model suggests a relationship to epileptic seizures and a resolution to a short-term memory controversy. Two EEG characteristics make other models implausible.

## 1. Introduction

This article is the fifth in a series of articles that show how neurons can be connected to process information. These articles show that the field of logic circuit design can inform neuroscience as well as vice versa. The first three articles [1–3] showed that a neural fuzzy logic decoder can produce the major phenomena of color vision and olfaction. The fourth article [4] showed that neurons can be connected to form Boolean neural flip-flops (NFFs) that are robust and generate the major phenomena of short-term memory. A flip-flop is a mechanism that can be set repeatedly to either one of two stable states, commonly labeled 0 and 1. Flip-flops are the basic building blocks of sequential logic systems, whose logic operations depend on both the current inputs and past sequence of inputs. Some of the material in [4] will be reviewed and used here.

The present article shows how neurons can be connected to generate major phenomena of electroencephalography. Only minimal properties of the strengths of excitatory and inhibitory signals are required for the networks to accomplish this. These properties are consistent with more complex neuron properties, such as synaptic plasticity and the effects of neuromodulators, but the minimal properties are sufficient to produce the results found here. The network proposed here together with NFFs proposed in [4] suggest a resolution to the longstanding controversy of whether short-term memory depends on neurons firing persistently or in brief, coordinated bursts [5, 6].

The hypothesis that cascaded oscillators produce the distribution of frequencies found in EEGs is consistent with available data, but further research is needed for a rigorous test. The hypothesis implies that the entire distribution of EEG frequencies in bands is determined by the mean and variance of the delay times of neurons that make up the initial oscillators in the cascades. With samples of neuron delay times and EEG frequencies, this implication can be tested simply and rigorously with standard statistical tests for equal means and variances.

The brain’s need for oscillations with the distribution of frequencies found in EEGs suggests a possible relationship between the oscillators proposed here and the abnormal electrical activity in neurological disorders such as epileptic seizures. This suggestion is summarized in the remainder of this introduction.

Oscillations in EEGs have a wide variety of frequencies, including high frequencies. From an engineering standpoint, connecting logic gates to produce oscillations with such a variety of frequencies is not a straightforward design problem. A toggle is a flip-flop with one input that inverts the state with each input pulse. Toggles connected in sequence are ideal for generating oscillations with a wide variety of frequencies because of the exponential growth in periods (doubling with each successive toggle). A three-inverter ring oscillator is the fastest oscillator that can be constructed with logic gates, and the distribution of frequencies that three-neuron ring oscillators can produce closely matches the distribution of frequencies in the EEG gamma band. But driving cascaded toggles with the ring oscillator presents several problems.

The master-slave toggle is the standard choice for cascaded toggles because a long input pulse inverts the toggle only once. A master-slave toggle can be constructed with neurons. But a three-neuron ring oscillator’s pulse duration is too short, and the oscillation frequency is too high, to invert a master-slave toggle correctly.

A JK flip-flop can be configured as a toggle, but it has a narrow range for an input pulse duration that can successfully invert it. The range is approximately three to four neuron delay times. Fortuitously, a three-neuron ring oscillator’s pulse duration is just within the upper bound.

The JK toggle’s output pulse duration is too long to invert another JK toggle correctly. Fortunately the toggle’s two initial neurons produce output pulses of nearly the same duration as the toggle input. One of these signals can be used to invert the next toggle in the cascade. The master-slave would work for subsequent toggles in the cascade after the first one, but since the simpler JK suffices, there may have been no selective pressure to find the master-slave. There are other possibilities for toggles besides the JK design given here, but they would also have the problem of a narrow requirement for input pulse duration.

Irregularities such as variations in neuron delay times could cause serious errors in the JK toggle because of its narrow margin for error. For example, if the delay times of the neurons in a cascade’s initial ring oscillator are substantially different from those in the cascade’s first toggle, that could cause an error. Delay times can be stabilized by averaging the signals from several neurons and by other error-correcting mechanisms such as neuromodulators. But with such a narrow margin for error, such methods may not be foolproof. Depending on the type of error that occurs, neural structures that are synchronized by the oscillator would either be disabled or enabled but unsynchronized. The resulting timing errors in neural firing and the brain’s efforts to deal with the errors could be related to the abnormal electrical activity characteristic of epileptic seizures.

The oscillator and JK toggle designs proposed here, as well as cascaded oscillators composed of JK toggles, are likely to be new to engineering.

## 2. Materials and methods

### 2.1. Simulation materials and methods

Simulations of electronic circuits were done in CircuitLab. A 5V signal represents the logic value TRUE, and 0V represents FALSE. The signal graphs are stacked for display by adding increments of 10V.

The neural oscillators and toggles were simulated in MS Excel. A neuron’s state is its output signal strength, normalized for convenience to be in the interval [0, 1]. For the simulations, the number t_i_ represents the time after i changes of state (i = 0, 1, 2, …). The time required for a neuron to change from one state to another is the neuron’s delay time. The neurons’ outputs are initialized at time t_0_ = 0. For i > 0, the output of each neuron at time t_i_ is computed as a function of the inputs at time t_i-1_. This function is given below. Baseline neuron activity and low level additive noise in neuron signals are simulated by a computer-generated random number uniformly distributed between 0 and 0.1.

Specific predicted probabilities of unusually high gamma band frequencies were approximated numerically from the estimated frequency probability density function (PDF) of the initial oscillator in the oscillator cascade with Converge 10.0, although this could also be done with a substitution of u = 1,000/x to convert the frequency PDF to a normal distribution of periods.

### 2.2. Unexplained phenomena and alternative models

#### 2.2.1. Short-term memory: persistent firing or brief, coordinated bursts?

Memory tests have shown that certain neurons fire continuously at a high frequency while information is held in short-term memory [7, 8]. These neurons exhibit seven characteristics associated with memory formation, retention, retrieval, termination, and errors. One of the neurons in the NFFs proposed in [4] was shown to produce all of the characteristics.

In addition to neurons firing persistently, other neurons firing in brief, coordinated bursts are also associated with short-term memory [5]. Which of these two phenomena actually produces short-term memory has been a longstanding controversy [5, 6]. It will be shown that neural oscillators and NFFs together suggest a resolution to this issue.

#### 2.2.2. Electroencephalography

##### 2.2.2.1. EEG phenomena and previous models

Electroencephalograms show widespread rhythms that consist of many neurons firing with matched periods. The spectrum of frequencies has been partitioned into bands according to the behavioral and mental state associated with the frequencies in each band. Five EEG frequency bands are considered here: gamma, beta, alpha, theta, and delta. Some researchers have found more bands or divided the bands into sub-bands depending on the focus of their research, but these five are discussed most often in the literature.

The distribution of frequencies within each of these bands is unimodal [9–12]. The ratios of consecutive band boundaries [13] and the ratios of consecutive band peak frequencies [9–12] are approximately 2. The gamma band peaks at about 40 Hz [9–12], although it contains frequencies of 100 Hz or more [14, 15]. Several estimated frequencies have been found for each boundary between bands.

The EEG phenomena raise several questions. What produces the widespread, synchronized, periodic firing? What is the function of this widespread synchronization? What produces and what is the function of the wide distribution of EEG frequencies in bands? What produces the unimodal distribution in each band and the octave relationships between the peaks and boundaries? What determines the specific frequencies of the peaks and boundaries? Why do gamma oscillations peak at about 40 Hz? Why does the gamma band contain frequencies that are considerably faster than 40 Hz? Why is there little agreement on the boundaries separating the EEG bands?

It will be shown that the cascaded oscillators model provides answers to all of the questions above.

Many models have been proposed for producing EEG frequencies [e.g., 16-23]. These models have several significant differences from the cascaded oscillators model. The models are far more complex than the cascaded oscillators’ minimal architecture (three or six neurons per oscillator) and neuron requirements (minimal capabilities of excitation and inhibition). The models are not explicit in the sense of showing all neurons and connections. Each model focuses on a narrow aspect of EEGs, such as how some frequencies can be generated in one or two frequency bands. The models do not produce any of the known characteristics of the distribution of EEG frequencies, such as the octave relationships between the bands or the specific band peaks or boundaries. The models also do not show that EEG phenomena can arise from selective pressures for a biologically useful function. None of the models can answer more than one or two of the questions above. And none has a micro-level explanation for its results like the mean and variance of neuron delay times that determine the entire distribution of frequencies produced by cascaded oscillators.

##### 2.2.2.2. Implausibility of alternative mechanisms

The cascaded oscillators design is based on evolutionary selective pressure for the biologically useful function of synchronization of neural structures’ state changes to avoid timing errors. This includes two specific selective pressures: for some types of information to be processed as fast as possible, and for a wide variety of speeds for diverse brain functions in the tradeoff between speed and accuracy. The first can be met by a three-inverter ring oscillator, the fastest oscillator that can be achieved by a network of logic gates. The second can be met by cascaded toggles, which double the period with each toggle.

Conceivably, the biological need for fast oscillations could be met by a mechanism that is different from a three-neuron ring oscillator, and a wide variety of speeds could be achieved by something other than cascaded toggles. But according to the available data for neuron delay times and EEG frequencies, the distribution of EEG gamma frequencies matches that of three-neuron ring oscillators. Also according to available data for EEG frequencies, the distributions of the other EEG bands halve the frequencies with each band. These two EEG features are by-products of a possible simple solution to selective pressures, but otherwise they have no apparent biological function. This makes it implausible that EEG phenomena are produced by a mechanism that is fundamentally different from cascaded oscillators.

### 2.3. Analysis

#### 2.3.1. Neuron signals

##### 2.3.1.1. Neuron signal strength

Neuron signal strength is normalized here by dividing it by the maximum possible strength for the given level of adaptation. This puts intensities in the interval from 0 to 1, with 0 meaning no signal and 1 meaning the maximum strength. The normalized number is called the *response strength* or simply the *response* of the neuron. The responses 1 and 0 are collectively referred to as binary signals and separately as high and low signals.

The strength of a signal consisting of action potentials, or spikes, is measured by spike frequency. A high signal consists of a burst of spikes at the maximum spiking rate. For a signal that oscillates between high and low, the frequency of the oscillation is the frequency of bursts (not to be confused with the frequency of spikes).

Normalization is only for convenience. Non-normalized signal strengths, with the highest and lowest values labeled Max and Min, rather than 1 and 0, would do as well. Absolute maximum and minimum values are also not necessary for high and low. These could be replaced by high and low ranges of values.

Table 1 shows a truth table for the logic function X AND NOT Y. The last column also represents the approximate response of a neuron with high and low excitatory and inhibitory inputs X and Y, respectively. Of the 16 possible binary functions of two variables, this table represents the only one that is consistent with the customary meanings of “excitation” and “inhibition.” In simplest terms, a neuron is active when it has excitatory input *and* does *not* have inhibitory input.

**Table 1.**
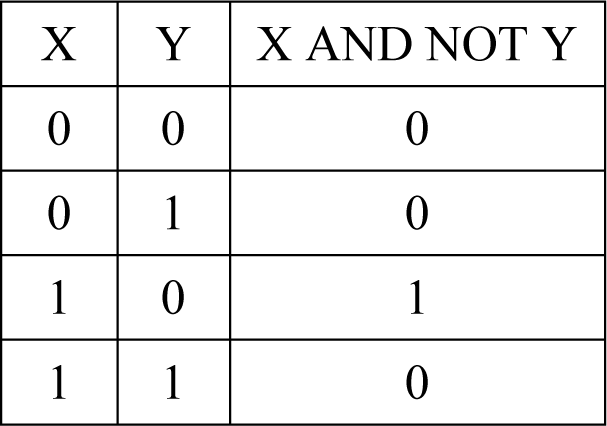
AND-NOT logic function. The table is a logic truth table for the statement X AND NOT Y. It also shows the approximate response of a neuron with one excitatory input of strength X and one inhibitory input of strength Y.

Some of the networks presented here require continuous, high input. In the figures, this input is represented by the logic value “TRUE.” For an electronic logic circuit, the high input is normally provided by the power supply. If the components represent neurons, the high input can be achieved by neurons in at least four ways. 1) A continuously high signal could be provided by a neuron that has excitatory inputs from many neurons that fire independently [24]. The brain has many neurons that are active spontaneously and continuously without excitatory input [25, 26]. A network neuron that requires a high excitatory input could receive it from 2) a spontaneously active neuron, or 3) the neuron itself could be spontaneously active. 4) The high input could be provided by one of an NFF’s outputs that is continuously high (or nearly high).

##### 2.3.1.2. Additive noise in neuron signals

###### 2.3.1.2.1 Noise reduction

A sigmoid response to excitatory input produces an output that is closer to binary than the input by decreasing an input near 0 and increasing an input near 1. Some neurons are known to have sigmoid responses to single inputs, including inhibitory inputs [27–29]. It will be demonstrated by simulation that a neuron response that is sigmoid in both excitatory and inhibitory inputs is sufficient to produce robust near-binary outputs of the NFFs and oscillators presented here. But a sigmoid response is not necessary; a simpler, more general property is sufficient.

Reduction of noise in both excitatory and inhibitory inputs can be achieved by a response function that generalizes a sigmoid function’s features. The noise reduction need only be slight for the proposed NFFs and oscillators because they have feedback loops that continuously reduce the effect of noise.

Let F(X, Y) represent a neuron’s response to an excitatory input with strength X and an inhibitory input with strength Y. The function value, as well as X and Y, must be bounded by 0 and 1, the normalized minimum and maximum possible neuron responses. Suppose F satisfies:

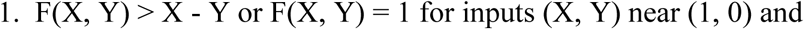

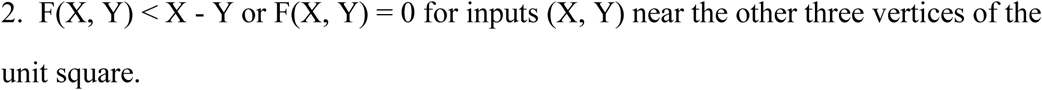

The truth values of the logic function X AND NOT Y in Table 1 are max{0, X-Y} (the largest of 0 and X-Y). For inputs X and Y that are somewhat close to binary, conditions 1 and 2 make the neuron’s response F(X, Y) closer to, or equal to, the binary truth value X AND NOT Y of Table 1 than max{0, X-Y}. Neurons that make up the networks proposed here are assumed to have these two minimal noise-reducing properties.

There are many examples of functions that satisfy conditions 1 and 2. A sigmoid neuron response to excitatory input with no inhibitory input satisfies the conditions. The simple “threshold” response function of a neuron that has no response for low inputs, and fires at a high rate when the excitatory input strength is above a certain threshold value, is an extreme form of a sigmoid response and satisfies the conditions. A function that is sigmoid in both excitatory and inhibitory inputs (e.g., the function in the next section) satisfies the conditions. Also shown below is a single transistor with two inputs that satisfies the conditions. Conditions 1 and 2 are also consistent with more complex models. For example, each of the two inputs X and Y could represent nonlinear weighted sums (or even more complex functions) of several excitatory and inhibitory inputs, respectively, and F could be a nonlinear weighted sum of X and Y.

Conditions 1 and 2 are sufficient to reduce additive noise in binary inputs and produce the NFF results found here. The level of noise that can be tolerated by the NFFs depends on the two regions in the unit square where condition 1 or 2 holds for the input point (X, Y). If a binary input (X, Y) has additive noise that is large enough to change the region in which it lies, an error can occur. Because of the continuous feedback in NFFs, it will be seen that even a response function with only a moderate noise-reducing capability can maintain near-binary outputs with substantial noise in the inputs.

###### 2.3.1.2.2. Example of a neuron response that satisfies conditions 1 and 2

For any sigmoid function f from f(0) = 0 to f(1) = 1, the following function has the noise-reducing properties 1 and 2:

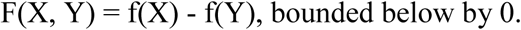

This function is plausible as an approximation of a neuron response because it is sigmoid in each variable and some neurons are known to have sigmoid responses to single excitatory or inhibitory inputs, as mentioned above. The same sigmoid function applied to X and Y is not necessary to satisfy conditions 1 and 2. The function F could be the difference of two different sigmoid functions.

The function F is illustrated in Fig 1 for a specific sigmoid function f. The sine function of Fig 1A, which is somewhat close to the line y = x, was chosen for f rather than any of the more common examples of sigmoid functions to demonstrate by simulation that a highly nonlinear function is not necessary for robust maintenance of binary signals. On half of the unit square, where Y ≥ X, Fig 1B shows that F has the value 0. This reflects the property that a large inhibitory input generally suppresses a smaller excitatory input.

**Fig 1.**
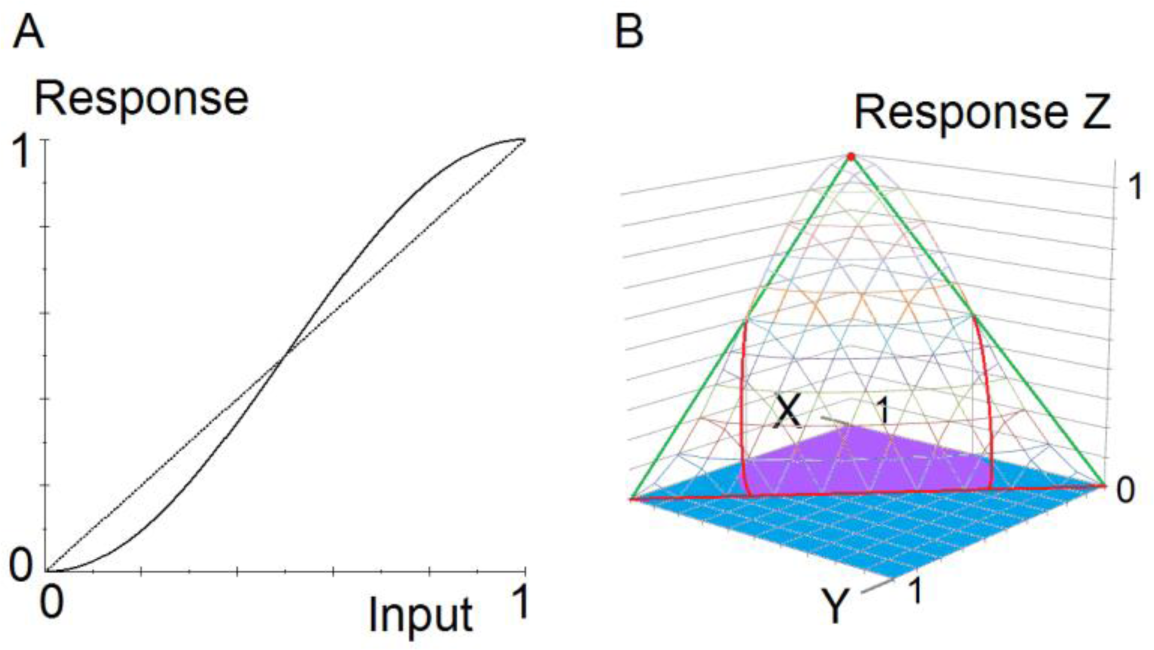
Noise-reducing function. The graphs show an example of a neuron response to analog inputs that reduces additive noise in binary inputs. **A.** A sigmoid function f(x) = (1/2)sin(π(x - 1/2)) + 1/2. **B.** Graph of F(X, Y) = f(X) - f(Y), bounded by 0. The response function has the noise-reducing properties 1 and 2. Wireframe: Graph of the function Z = F(X, Y). Green and red lines: A triangle in the plane Z = X - Y. Red: Approximate intersection of the plane and the graph of F. Purple: Approximate region in the unit square where F(X, Y) > X - Y or F(X, Y) = 1 (condition 1). Blue: Approximate region in the unit square where F(X, Y) < X - Y or F(X, Y) = 0 (condition 2).

###### 2.3.1.2.3. Neuron response for simulations

The response function F(X, Y) in Fig 1 is used for the simulations as follows. The number t_i_ represents the time after i changes of state. The increment t_i_ - t_i-1_ is the delay time of the neuron that changed states. The neurons’ outputs are initialized at time t_0_ = 0. At time t_i_ for i > 0, the output Z_i_ of each neuron that has excitatory and inhibitory inputs X_i-1_ and Y_i-1_ at time t_i-1_ is:

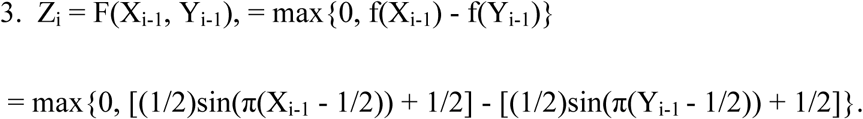

###### 2.3.1.2.4. A primitive noise-reducing gate

Properties 1 and 2 do not indicate sophisticated capabilities of mathematics or logic. A response with the properties can be produced by mechanisms that are quite simple. Fig 2 shows that a single transistor and three resistors can be configured to accomplish this. The inputs X and Y vary from 0V to 5V in steps of 0.05V.

**Fig 2.**
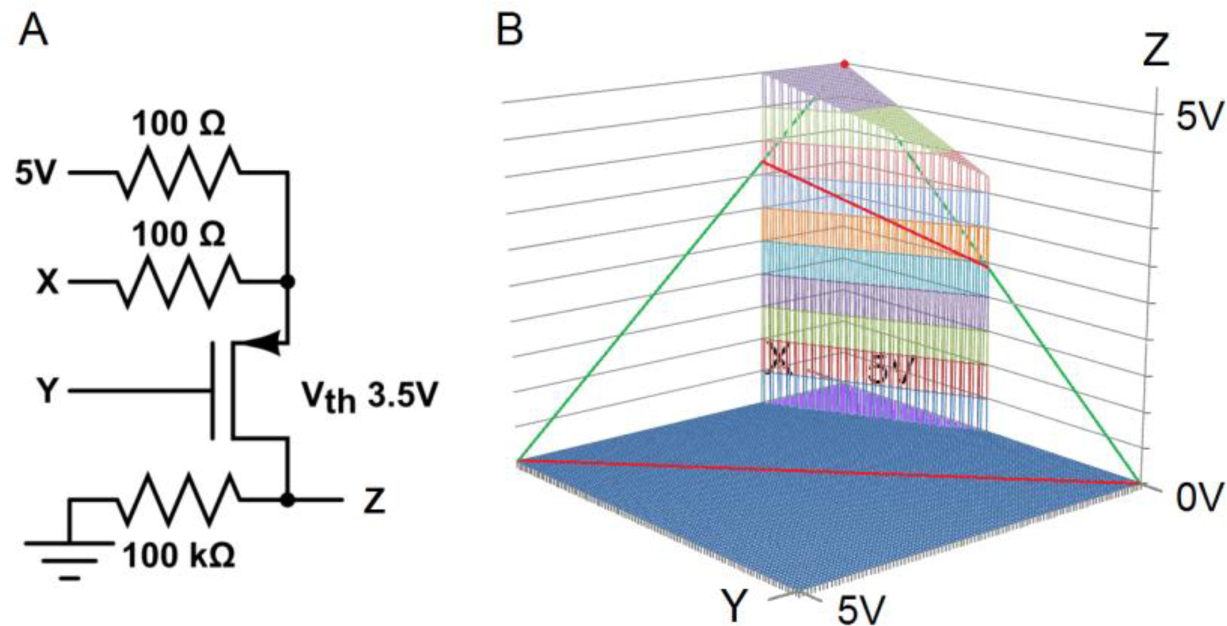
Single transistor gate that reduces noise. This minimal logic circuit satisfies the noise-reducing conditions 1 and 2. **A.** A logic circuit consisting of one transistor and three resistors. **B.** CircuitLab simulation of the logic circuit in A. Wireframe: Graph of the transistor response function Z = F(X, Y). Three green and red lines: A triangle in the plane Z = X - Y. Red: Intersection of the plane and the graph of F. Purple: Region in the unit square where F(X, Y) > X - Y or F(X, Y) = 1 (condition 1). Blue: Region in the unit square where F(X, Y) < X - Y or F(X, Y) = 0 (condition 2).

#### 2.3.2. Neural logic gates and flip-flops

For several reasons that were detailed in [4], the neural networks in the figures are illustrated with standard (ANSI/IEEE) logic symbols rather than symbols commonly used in neuroscience schematic diagrams. One of the reasons is that the symbols can be interpreted in two ways. As a logic symbol, the rectangle with one rounded side in Fig 3A represents the AND logic function, and the circle represents negation. The input variables X and Y represent truth values TRUE or FALSE, and the output represents the truth value X AND NOT Y. Second, Fig 3A can also represent a single neuron, with a circle representing inhibitory input and no circle representing excitatory input. With the minimal noise-reducing capabilities of conditions 1 and 2, if X and Y are binary with some additive noise, the output will be closer to, or equal to, the binary X AND NOT Y value of Table 1 than max{0, X-Y}.

**Fig 3.**
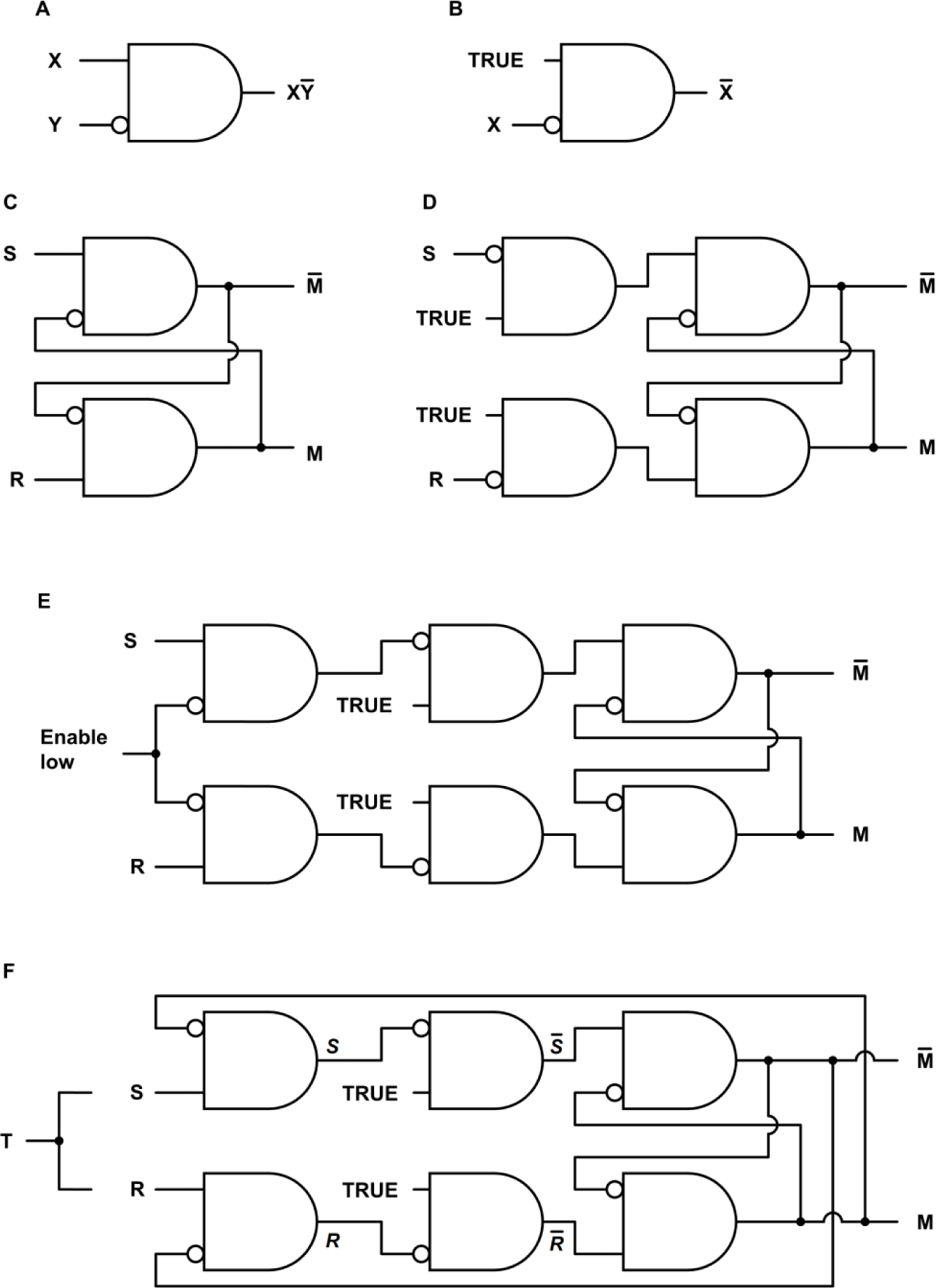
Neural logic gates and flip-flops. **A.** A symbol for an AND-NOT logic gate, with output X AND NOT Y. The symbol can also represent a neuron with one excitatory input and one inhibitory input. **B.** An AND-NOT gate configured as a NOT gate, or inverter. **C.** An active low Set-Reset (SR) flip-flop. **D.** An active high SR flip-flop. **E.** An active high SR flip-flop enabled by input from an oscillator. **F.** A JK flip-flop or toggle.

As logic circuits, the rest of the figures’ outputs shown in Fig 3 follow from Boolean logic. The figures’ outputs as networks composed of neurons will be illustrated as needed by simulation. Fig 3B shows an AND-NOT gate with a continuously high input that functions as a NOT gate, or inverter.

A flip-flop stores a discrete bit of information in an output with values usually labeled 0 and 1. This output variable is labeled M in Fig 3. The value of M is the flip-flop *state* or *memory bit*. The information is stored by means of a brief input signal that activates or inactivates the memory bit. Input S *sets* the state to M = 1, and R *resets* it to M = 0. Continuous feedback maintains a stable state. A change in the state *inverts* the state.

Two basic types of flip-flops are the Set-Reset (SR) and JK. Fig 3C shows an active low SR flip-flop. The S and R inputs are normally high. A brief low input S sets the memory bit M to 1, and a brief low input R resets it to 0. Adding inverters to the inputs in Fig 3C produces the active high SR flip-flop of Fig 3D. The S and R inputs are normally low. A brief high input S sets the memory bit M to 1, and a brief high input R resets it to 0. A disadvantage of the SR flip-flop is that if S and R input signals are attempting to invert the flip-flop simultaneously, the outputs are unpredictable.

Fig 3E shows a flip-flop with an enabling input. The S and R inputs in Fig 3D have been replaced by AND-NOT gates that allow the S or R input to be transmitted only when the enabling input is low. In synchronized signaling systems, several logic circuits are enabled by an oscillator signal to avoid timing errors. Structures that are enabled by the same signal change states simultaneously. Adding an inverter (Fig 3B) to the enabling signal can ensure that one structure does not change states simultaneously with another.

For the so-called JK flip-flop in Fig 3F, the enabling input in Fig 3E has been replaced by input from the flip-flop outputs. The advantage of the JK flip-flop over the SR flip-flop is that if S and R are both high simultaneously, the flip-flop state is inverted because one of the two input gates is inhibited by one of the flip-flop outputs. This means the JK flip-flop can be configured as a toggle by linking the Set and Reset inputs, as illustrated by the single input T in the figure.

A problem with the JK toggle is that it functions correctly only for a short duration of high input. If the input pulse is too long or too short, an error occurs. This problem is discussed in the toggle section below.

#### 2.3.3. Neural toggles and oscillators

##### 2.3.3.1. Neural toggle

Fig 4 shows a simulation of the JK flip-flop of Fig 3F composed of neurons and configured as a toggle. The outputs were initialized in a stable state, and the simulation was carried out as described in the simulation methods with equation 3. The slow rise and fall of the toggle input T, over several state changes, is exaggerated to clarify the robust operation of the network in the presence of additive noise.

**Fig 4.**
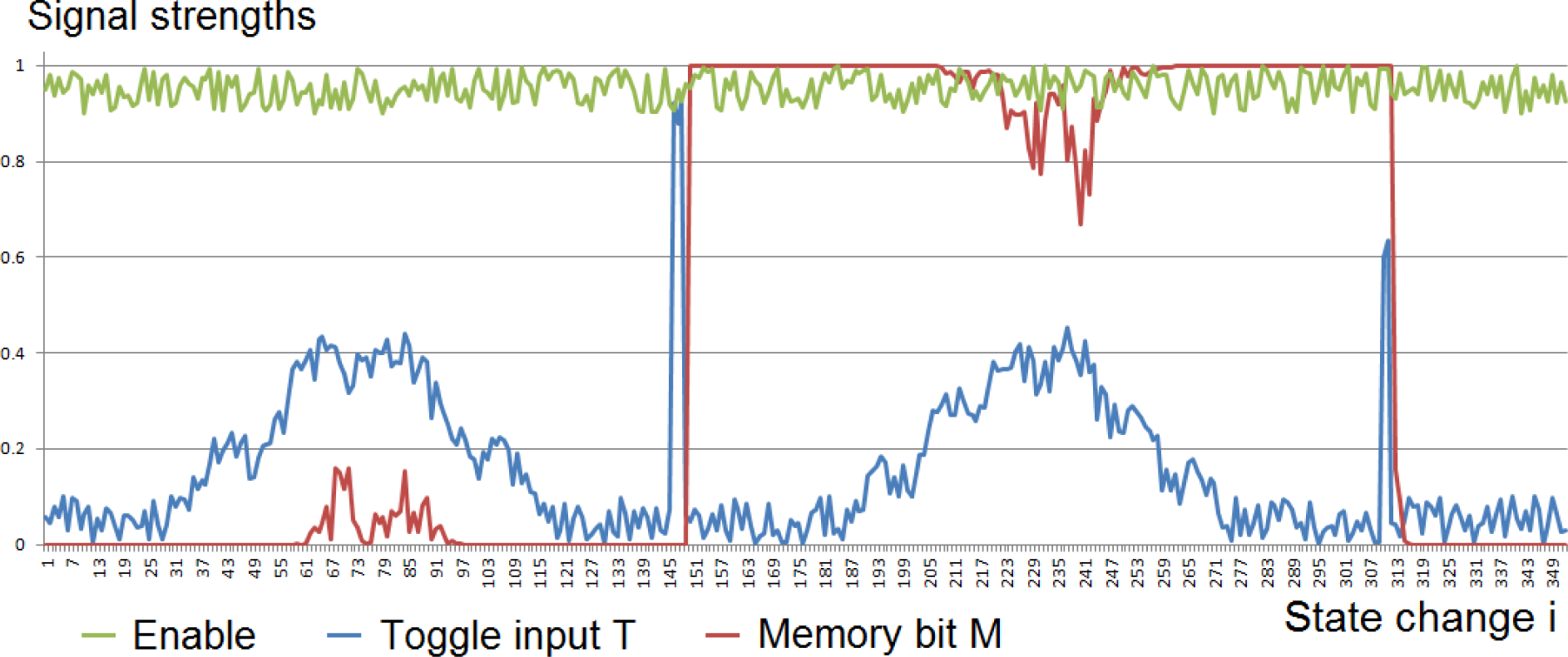
Simulation of a neural toggle with noise in the inputs. The figure shows a simulation of the JK flip-flop in Fig 3F composed of six neurons and configured as a toggle. The input TRUE in Fig 3F is labeled Enable here because it contains additive noise. The graphs show the toggle’s operation is robust in maintaining a nearly binary memory bit in the presence of substantial additive noise in the inputs. Baseline noise in both the Enable input and the toggle input T has negligible effect on the memory bit M. Two temporary bursts of larger noise in T have no lasting effect. The two inputs that invert the toggle state have strengths 0.85 + noise and 0.55 + noise, with durations of four and three state changes, respectively. The values 0.85 and 0.55 were chosen to show that high inputs subtantially reduced by noise as well as high inputs successfully invert the toggle.

Low level additive noise and baseline activity in the inputs in Fig 4 are simulated by a computer-generated random number uniformly distributed between 0 and 0.1. The high Enabling input is simulated by 1 minus noise. Each of the two temporary bursts of larger noise in T is simulated by the sum of two sine functions and the computer-generated noise.

The JK toggle is sensitive to the duration of the high input pulse T. The simulations in Fig 5 illustrate the limitations on the input pulse durations. The simulations were carried out as described in the simulation methods with equation 3.

**Fig 5.**
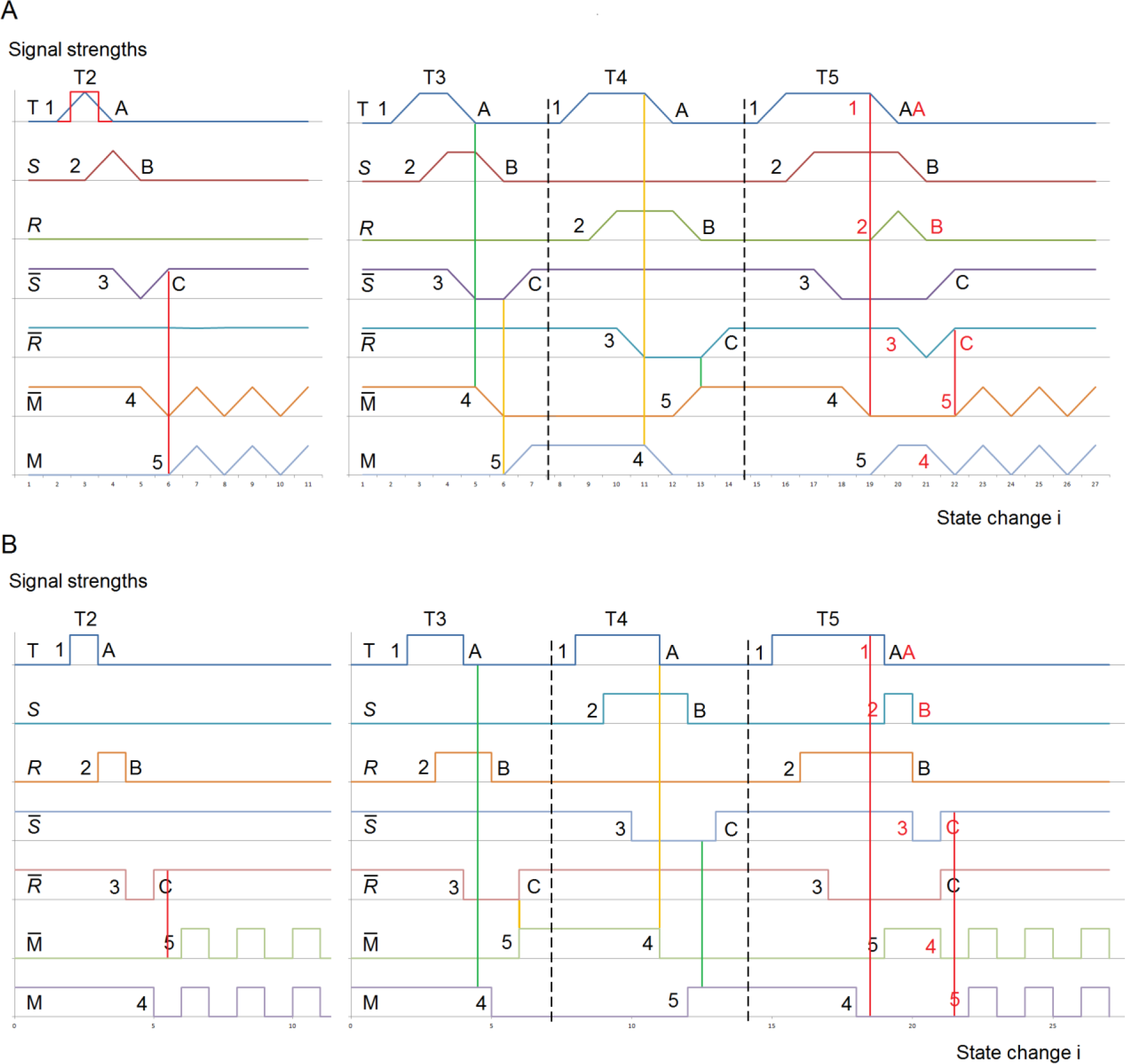
Neural toggle simulations illustrating the narrow requirement for input pulse duration. The graphs show the simulated responses of a neuron implementation of the JK toggle in Fig 3F, with high input pulses T that last for two to five neuron changes of state. These pulse durations include the pulse’s rise and fall. The piecewise linear (PWL) graphs in simulation **A** and piecewise square (PWS) graphs in simulation **B** indicate rise and fall that are spread uniformly over the change of state or that occur quickly, respectively. For comparison, the PWS pulse T2 from Fig 5B is superimposed in red on the PWL T2 in Fig 5A. The results in the two simulations are the same, except that wherever one illustrates a Set (*S*) inversion, the other shows a Reset (*R*).

A successful inversion of the toggle state requires the input T to be high long enough to invert the toggle, but not so long that it initiates a second toggle inversion. An input pulse that is too short or too long causes the inhibiting feedback from a toggle output to begin too late or to end too soon, respectively. For each high pulse of the toggle input T, the sequence of signal inversions numbered 1 through 5 in Fig 5 inverts the two toggle outputs. The sequence A, B, C terminates the toggle inversion.

If the input pulse T is too short, the terminating sequence A-C completes before the inverting sequence 1-5. That is, step C occurs before step 5, which means the inhibiting feedback to the low toggle output begins too late to prevent an extra inversion of that output. This error occurs for T2 in Fig 5, highlighted with a red line. In Fig 5A for example, step 5 makes M high, which is supposed to keep M_bar inhibited in a low state. But because step C returned *S*_bar to the high state before step 5 provides the inhibitory high input from M, M_bar returns to the high state.

In the simulation, the toggle outputs M and M_bar happen to go high simultaneously and they continue oscillating together in a race condition. As will be seen, real materials do not make such changes exactly simultaneously. One of the two outputs will soon win the race, stabilizing the outputs.

The too-short input error is only narrowly avoided in the case T3, where steps C and 5 occur in the same change of state. The near error is highlighted with a yellow line. In case T4, step C occurs after step 5, highlighted with a green line.

If the input pulse T is too long, the high toggle output inverts too soon to inhibit the toggle input from initiating another toggle inversion. That is, step 4 occurs before step A. This error occurs for T5 in Fig 5, highlighted with a red line. In Fig 5A for example, the toggle input T is still high when M_bar has inverted to low and is no longer suppressing *R*. This begins another inversion of the toggle state, shown in red. As before for short inputs, step C occurs before step 5, resulting in the race condition.

The too-long input error is narrowly avoided in the case T4, where steps 4 and A occur in the same change of state. The near error is highlighted with a yellow line. In case T3, step 4 occurs after step A, highlighted with a green line. Longer pulses in T simply invert the toggle twice, leaving the toggle state unchanged.

The next simulation may be more accurate in how real components actually function. Fig 6 shows an electronic implementation of an AND-NOT gate and two simulations of the JK toggle of Fig 3F composed of these gates. The threshold for each transistor is 2V.

**Fig 6.**
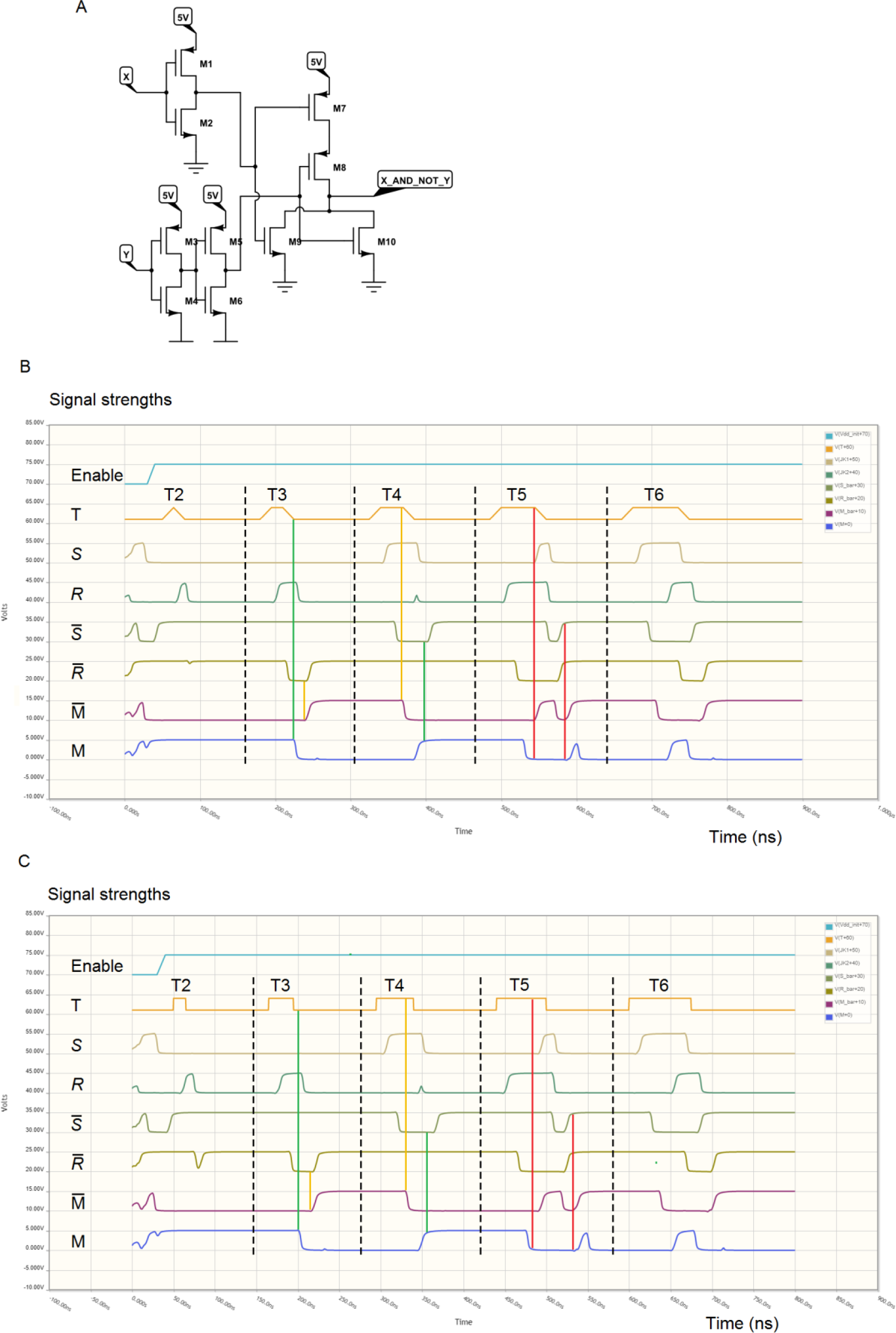
An electronic simulation that verifies the narrow requirement for the JK toggle’s input pulse duration. **A.** A CMOS AND-NOT gate of Fig 3A composed of a NOR gate (NOT OR), an inverter for X, and a buffer for Y to even the timing of the inputs. By De Morgan’s law, the output NOT[NOT(X) OR Y] = X AND NOT Y. **B.** A simulation of the JK toggle in Fig 3F composed of the AND-NOT gates in A with PWL inputs. The simulation time is 900 ns. **C.** A simulation of the toggle with PWS inputs. The simulation time is 800 ns.

The next electronic simulation will show that the delay time of the AND-NOT gate in Fig 6A is about 15 ns. So the input pulse durations for T1-T5 in Figs 6B and 6C are multiples of 15 ns. The electronic simulation software initializes the component states, so only one enabling input is needed for an asymmetry: the first 5V input to the first AND-NOT gate of the toggle.

Except for the race conditions and time scales, the simulations of the electronic toggle in Fig 6 show results that are similar to the simulations of the neural toggle in Fig 5. In Fig 6, the pulse T2 is simply too short to invert the toggle outputs. Pulses T3 and T4 invert the toggle outputs correctly. For the near error in T4, highlighted with yellow lines in Figs 6B and 6C, the pulse duration is long enough to produce a slight rise in *R,* the beginning of another toggle inversion. But the rise is too small to have a discernable effect on subsequent steps. Input T5 successfully inverts the outputs, but the long input is enough to raise a brief pulse in *S*. This causes the toggle outputs to be briefly malformed. Input T6 is so long it inverts the toggle twice, ultimately leaving the outputs unchanged. Longer input pulses also invert the toggle outputs only twice. Fig 6C shows a somewhat greater response to the T2 input than Fig 6B, especially evident in *R*_bar, because the high portion in the PWS input pulse T2 is wider than in the PWL pulse. The change in *R*_bar is not enough to have a ‘discernable effect on M.

The AND-NOT gate of Fig 6A was used for the simulations because of the highly reliable CMOS architecture. Although one-transistor AND-NOT gates like the one in Fig 2A can form functioning logic circuits [30, 31], the capability of electronic signals to travel both ways between components can cause problems in circuits composed of such simple hardware. Neural synapses have the distinct advantage of conveying signals that are generally transmitted only one way. This signal difference is one of the reasons that CMOS architecture is used almost exclusively in electronic logic circuits, and the reason that the AND-NOT gate in Fig 6A requires 10 transistors to accomplish what a single neuron can do.

The electronic simulations allow much more granular data than the neural simulations using equation 3, which evaluates only at the end of each state change. The simulation of Fig 6B was done with time steps of 1 ns, i.e., 15 evaluations for each state change. For Fig 6C, the time steps were 0.5 ns to simulate the PWS input more accurately.

##### 2.3.3.2. Neural ring oscillator

An oscillator produces periodic bursts of a high signal followed a quiescent period of a low signal. It is the basic element of a timing mechanism. A ring oscillator is a simple, reliable oscillator. An odd number of three or more inverters connected sequentially in a ring produces periodic bursts as each gate inverts the next one. The odd number of inverters makes all of the inverter states unstable, so the states oscillate between high and low. All inverters in the ring produce oscillations with the same frequency. Their phases are approximately uniformly distributed over one cycle, and the high and low signal durations are approximately equal. Their common period is twice the sum of the inverters’ delay times. (The sum is doubled because each component inverts twice per cycle.) A ring oscillator is the simplest type of oscillator that can be implemented with logic gates, and the simplest and fastest ring oscillator consists of three inverters.

Fig 7 shows a three-inverter ring oscillator composed of AND-NOT gates and two simulations of the oscillator composed of three neurons. Enable 1 rises two state changes before Enable 2 to initialize the cells. Additive noise in the Enable inputs is simulated by a random number uniformly distributed between 0 and 0.1. The Enabling input begins as baseline noise and transitions between 0 and 1 as a sine function plus noise. During the interval when the Enabling input is high, it is 1 minus noise.

**Fig 7.**
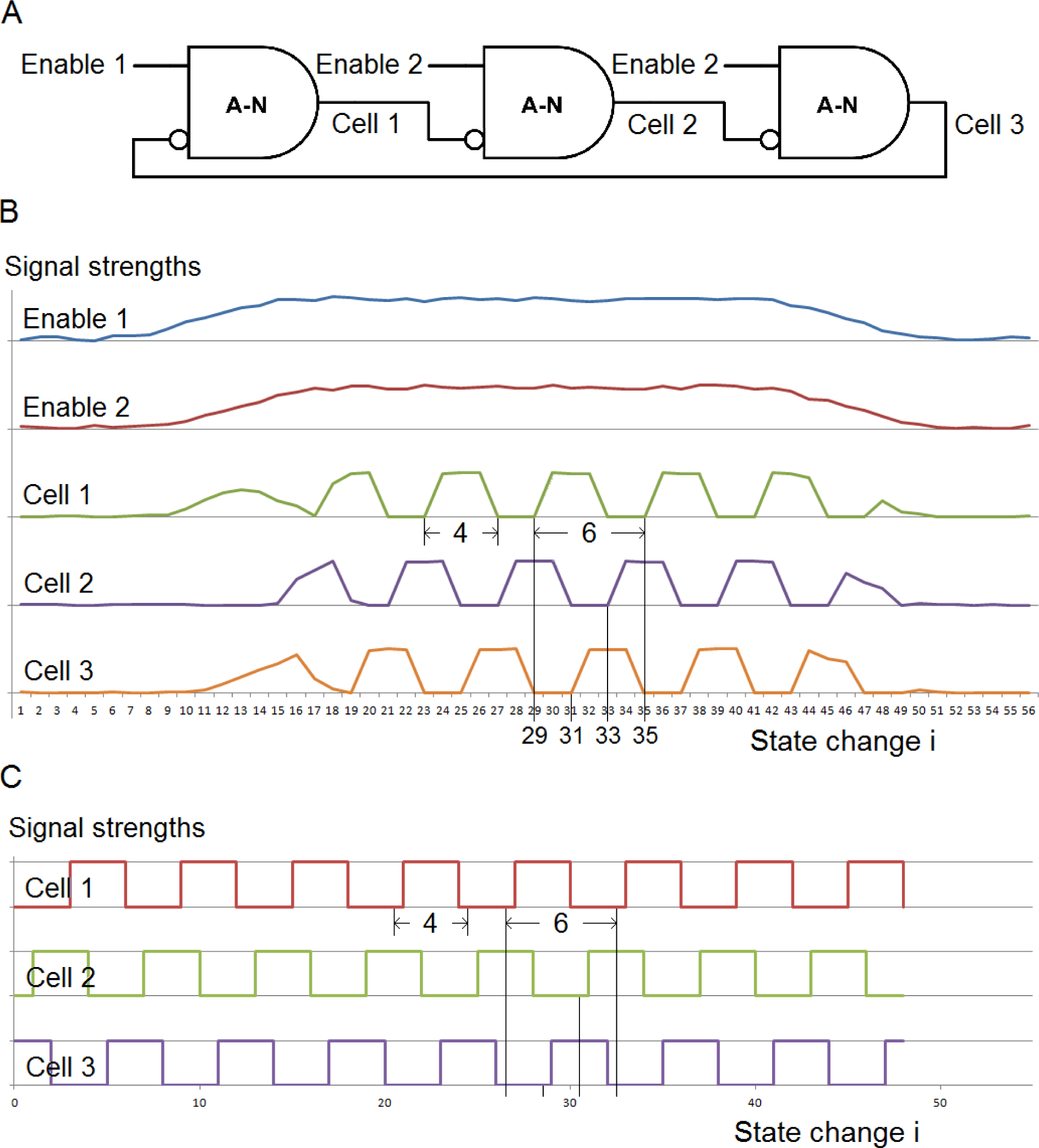
Ring oscillator. **A.** A ring oscillator consisting of three inverters of Fig 3B. **B.** A simulation of the ring oscillator composed of three PWL neurons. **C.** A simulation of the ring oscillator composed of three PWS neurons. The common period of six changes of state is the sum of the neurons’ delay times (two changes of state for each neuron). The phases are approximately uniformly distributed over the period, indicated by the black lines. The duration of each neuron’s high burst is four changes of state. This is a burst duration that can successfully invert a JK toggle.

Fig 8 shows a simulation of an electronic implementation of the three-inverter oscillator for comparison with the neural implementation. The simulation time is 650 ns.

**Fig 8.**
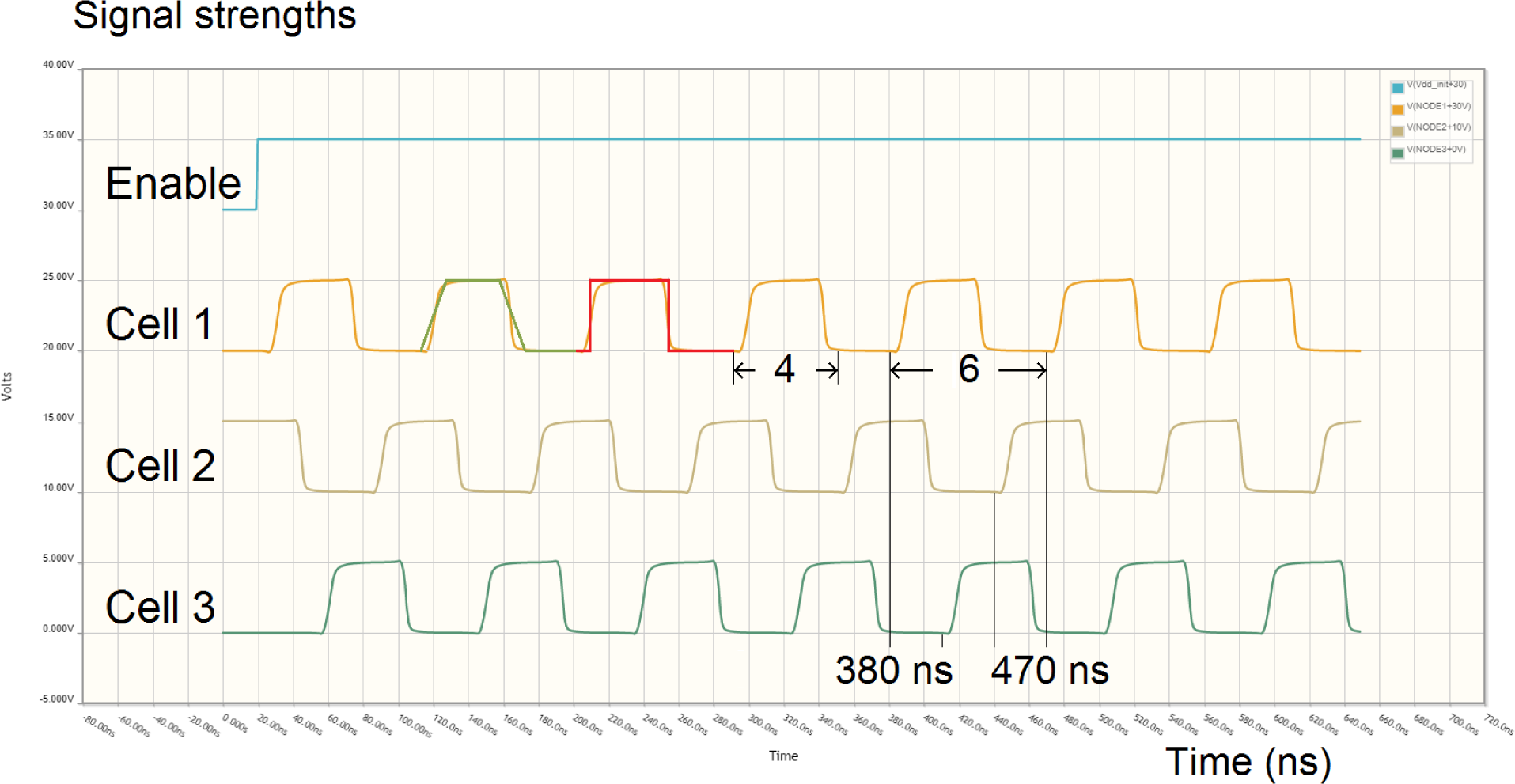
Simulation of an electronic ring oscillator. The graphs show a simulation of the oscillator in Fig 7A composed of AND-NOT gates in Fig 6A. Except for the time scale, the electronic simulation has essentially the same results as the neural simulations in Fig 7. One cycle of the PWL wave of Fig 7B and one cycle of the PWS wave of Fig 7C are superimposed for comparison. The slopes of the electronic oscillator’s rise and fall lie between the two extremes of the simulated neural oscillators’ PWL and PWS waves. The period of about 90 ns means the AND-NOT gate’s simulated delay time is about 15 ns. The three phases are approximately uniformly distributed over one cycle, as indicated by the black lines.

##### 2.3.3.3. Cascaded neural oscillators

As described in the introduction, an oscillator can be connected in sequence with toggles to form a cascade of oscillators. Because two high inputs are required for each cycle of a toggle-as-oscillator (one to set the memory state, another to reset it), a toggle produces a signal whose period is exactly double that of the toggle’s input. Fig 9 shows a cascade consisting of a ring oscillator and two toggles.

**Fig 9.**
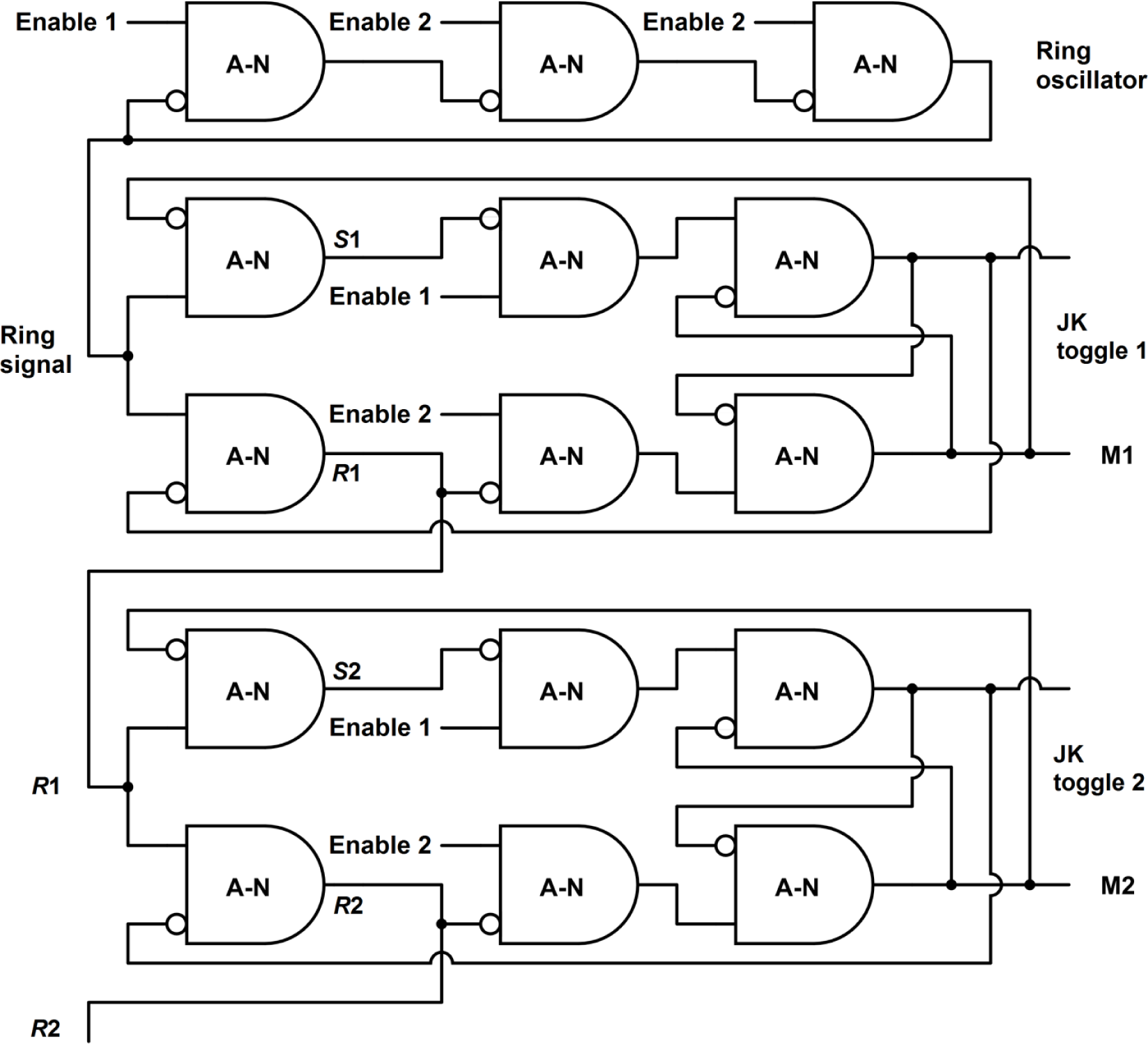
Three cascaded neural oscillators. The cascade consists of the ring oscillator of Fig 7A and two JK toggles of Fig 3F connected in sequence. A cascade could have any number of toggles. The input to the first toggle comes from the ring oscillator, and the input to each succeeding toggle comes from one of the first gates in the previous toggle. This makes the duration of each toggle’s high input pulse remain approximately the same throughout the cascade.

Simulations of the cascaded oscillators in Fig 9 are shown in Fig 10. Except for the time scales, the simulated neural results in Figs 10A and 10B are essentially the same as the simulated electronic results of Fig 10C.

**Fig 10.**
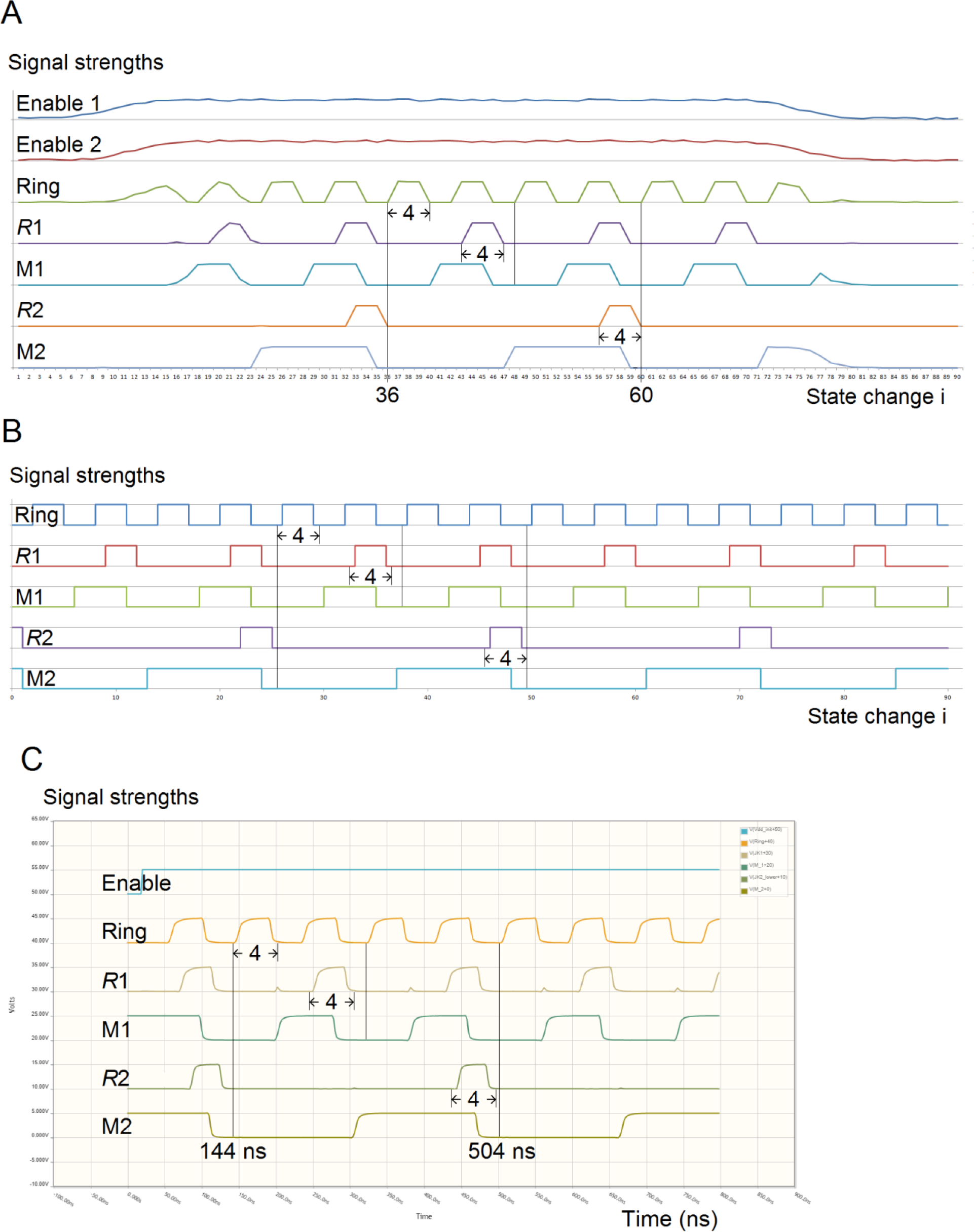
Simulations of the three cascaded oscillators in Fig 9. **A.** Simulation of oscillators composed of PWL neurons. **B.** Simulation of oscillators composed of PWS neurons. **C.** Simulation of oscillators composed of the electronic AND-NOT gates of Fig 6A. The simulation time is 800 ns.

The simulations in Fig 10 illustrate the main properties of cascaded oscillators: In each toggle, the period of every neuron’s output is double the period of the toggle’s input, as indicated by the black lines. The pulse duration of each toggle’s two initial gates is approximately the same as the pulse duration of the toggle’s input, as indicated by the measures of four changes of state. The duration of each toggle’s output pulse is approximately half of the period.

Because the input Ring to the first toggle has a pulse duration of four AND-NOT gate state changes, signal *R*1 in Fig 10C has a small rise in each cycle similar to the rise in *R* for input T4 in Fig 6, indicating the near error of the long input. Signal *R*2 in Fig 10C also has a rise at the same place in each cycle, but it is barely visible because the input *R*1 to the second toggle has a slightly shorter duration than the input Ring to the first toggle. Similarly, the pulse duration of *R*2 is slightly shorter than that of *R*1. If neurons have this behavior of a slight shortening of each input pulse to consecutive toggles in the cascade, it makes the cascade more robust because a pulse duration of four state changes is near the upper bound on pulse durations that can successfully invert the JK toggle.

#### 2.3.4. The relationship between the distributions of neuron delay times and cascaded neural oscillators’ frequencies

The distributions of cascaded neural oscillators’ frequencies are determined by the mean and variance of neuron delay times of the cascades’ initial oscillators.

##### 2.3.4.1. Exact relationships between inverter delay times and cascaded oscillator frequencies

The interest here is in neural inverters and toggles, but the arguments in this section apply to any implementation of inverters and toggles, including electronic. These results may not be found in electronics texts because engineers are not normally concerned with the small variances in component performance.

###### 2.3.4.1.1. Distributions of oscillator periods and frequencies

As noted earlier, each cycle time of a ring oscillator is the sum of the times it takes for each inverter to invert twice. If X_1_, …, X_n_ are independent and identically distributed random variables representing the delay times of n inverters in a ring oscillator, the ring oscillator’s period is:

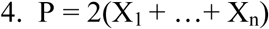

If toggles are connected in sequence with the oscillator, each cycle time of each toggle’s output is the sum of two of the input’s cycle times. Cascaded toggle number k = 1, 2, … has period:

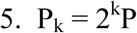

The mean and standard deviation of the delay times of the inverters in ring oscillators in all cascaded toggles are denoted by μ_d_ and σ_d_. By equations 4 and 5 and the elementary properties of random variables, for i = 1, 2, … (with i = 1 representing the initial ring oscillator), the period of cascaded oscillator number i has mean and standard deviation:

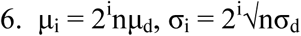

The factor 2^i^ shows the octave relationship between the oscillators’ distributions of periods.

The oscillators’ distributions of frequencies can be derived from the distributions of periods by straightforward calculus. If periods and frequencies are measured in milliseconds and hertz, respectively, then frequency = C/period for C = 1,000. If the probability density function (PDF) of the period of oscillator i = 1, 2, … (with i = 1 representing the initial ring oscillator) is f_i_(x), then the PDF of the frequency of oscillator i is:

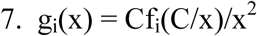

Equation 7 shows the oscillator period and frequency distributions are different. For example, it will be seen that if the periods are normally distributed, the frequency distributions are skewed to the right. But the intersections of consecutive period PDFs (converted to frequencies) are the same as the intersections of consecutive frequency PDFs because x^2^ and the initial constant C in equation 7 drop out of the equation g_i_(x) = g_i+1_(x).

###### 2.3.4.1.2. Normal distributions

If inverter delay times are normally distributed, then by equations 4 and 5 and the elementary properties of normal distributions, the periods of ring oscillators and cascaded toggles are also normally distributed.

The normal PDF with mean μ and standard deviation σ, whose graph is commonly known as the bell curve, is:

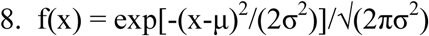

Equation 8 implies that a normal distribution is entirely determined by its mean and standard deviation. By equations 6 and 7, this means cascaded oscillators’ distributions of periods and frequencies are entirely determined by the number of inverters n in the initial ring oscillators and the inverter delay parameters μ_d_ and σ_d_.

Substituting the cascaded oscillators’ period parameters in equation 6 into equation 8 to obtain the period PDFs f_i_(x), the intersections of each pair of consecutive period PDFs can be found by elementary algebra. For i = 1, 2, …, (i = 1 representing the ring oscillator), the intersection of f_i_(x) and f_i+1_(x) occurs at period:

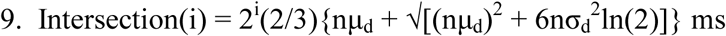

The factor 2^i^ shows the intersections also have the octave relationship.

By substituting the period PDFs f_i_ obtained from equations 6 and 8 into equation 7, the peak frequency (mode) for PDF g_i_ can be found by calculus:

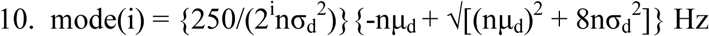

Again, the factor 2^i^ shows the peak frequencies also have the octave property. These peak frequencies are close to, but not the same as, the peak frequencies 1,000/μ_i_ derived from the means μ_i_ of the period’s normal distributions in equation 6.

##### 2.3.4.2. Neuron delay times

Since neuron delay times are determined by several factors, the delay times are approximately normally distributed (by the central limit theorem). For small networks with chemical synapses, nearly all of the delay occurs at the synapses. Several studies have measured synapse delay times [e.g., 32, 33], but the literature apparently does not have empirical estimates of the parameters (mean and variance) of the delay times’ distribution. However, a description of the range of synapse delay times is “at least 0.3 ms, usually 1 to 5 ms or longer” [25]. Although the description is far from precise, delay time parameters can be estimated.

The description of the range has two parts. The first part “at least 0.3 ms” seems to refer to all observations. The second part “usually 1 to 5 ms or longer” seems to describe the ranges of typical samples, with “5 ms or longer” representing the ranges’ right endpoints. In that case, the interval [1 ms, 7 ms] is at least a reasonable, rough estimate of the range of a moderately sized sample.

If only the range of a sample (minimum value, m, and maximum, M) is known, the midpoint can be used as an estimate of the mean of a distribution. Simulations have shown that (M - m)/4 is the best estimator of the standard deviation for moderately sized samples [34]. Based on this and the estimated range [1 ms, 7 ms], neuron delay times are estimated to have distribution parameters:

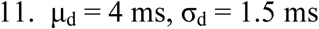

For a normal distribution with these parameters, about 99.3% of the distribution is at least 0.3 ms. This agrees well with the description “at least 0.3 ms.” About 73% lies between 1 and 5 ms, and 95% is between 1 and 7 ms. This agrees reasonably well with the description “usually 1 to 5 ms or longer.”

##### 2.3.4.3. EEG frequency distribution compared to estimated neural oscillator frequency distributions

The graphs of the estimated frequency PDFs of five cascaded neural oscillators are shown in Fig 11. As before, the period PDFs f_i_(x) are obtained by substituting the period parameters in equation 6 into equation 8. With the estimated delay parameters of equations 11 and n = 3 neurons for the ring oscillator, the estimated frequency PDFs g_i_(x) are obtained from equation 7. The four intersections of consecutive PDFs, shown in blue, are found by converting the periods given by equation 9 to frequencies. The five PDF modes, shown in yellow, are obtained from equation 10. Frequencies that are commonly cited [9-12, 35-52] as partition points separating the EEG frequency bands and peak frequencies of three of the bands are shown in red and green, respectively. Numbers in parentheses show how many times each frequency was cited. (Estimates of peak frequencies apparently have not been found for the lower frequency delta and theta bands.)

**Fig 11.**
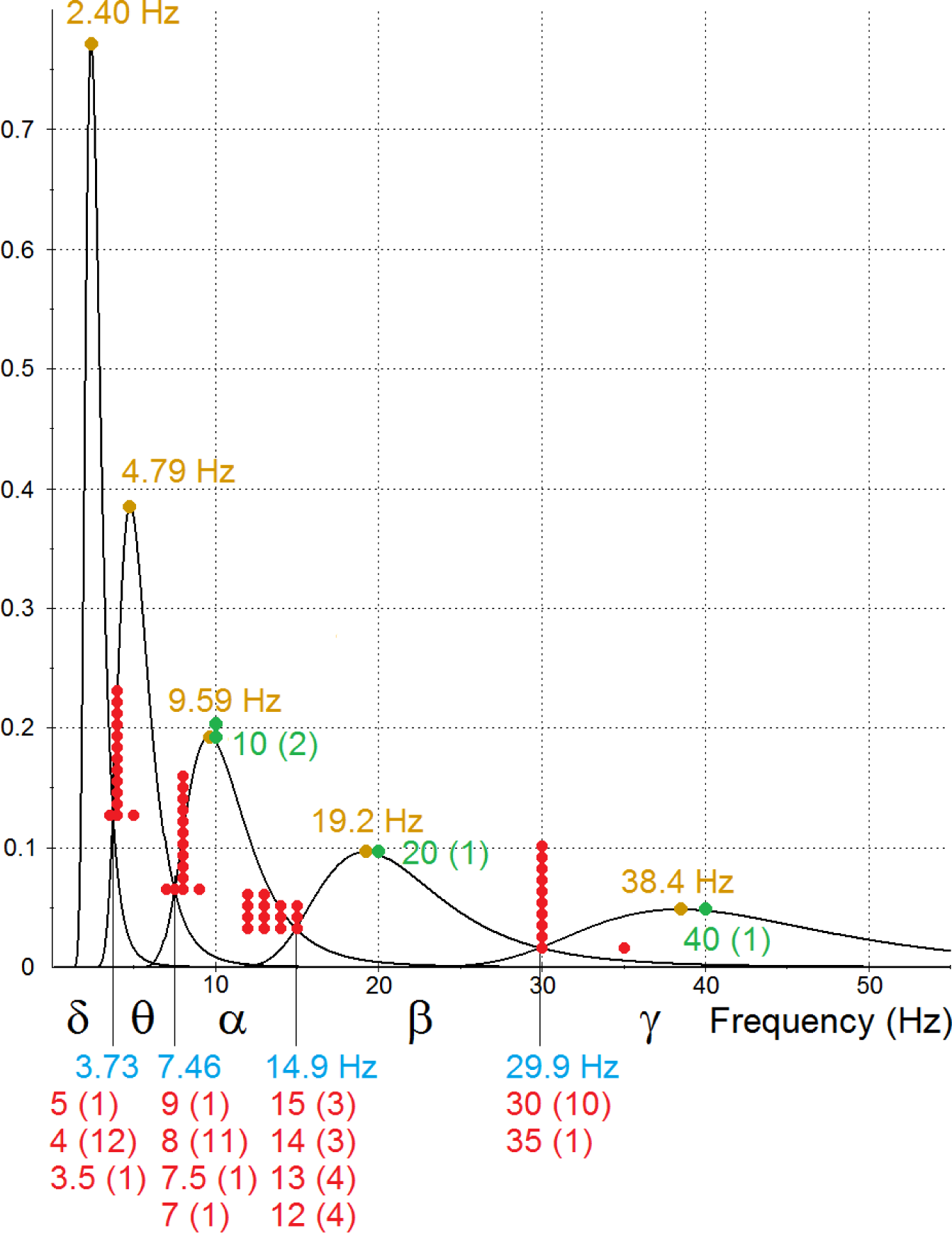
Estimated frequency distributions of cascaded neural oscillators compared to commonly cited EEG frequency band peaks and boundaries. The graphs are the estimated PDFs of the frequencies of a three-neuron ring oscillator and four cascaded toggles. The PDFs were determined solely by the estimated mean and variance of neuron delay times. The five intervals defined by the intersections of consecutive PDFs are labeled with Greek letters to distinguish them from EEG frequency bands, which are often written in the Roman alphabet. The intersections and modes are labeled in blue and yellow, respectively. Also shown in red and green are frequencies that are commonly cited as partition points separating the EEG frequency bands and peak frequencies of three of the bands. Numbers in parentheses and numbers of data points show how many times each frequency was found to be cited in a literature search. As predicted by the cascaded oscillators hypothesis, the graphs show that the modes and intersections of the estimated oscillator frequency PDFs are close to the peaks and partition points commonly cited for the EEG frequency bands, respectively.

#### 2.3.5. Synchronization

The EEG frequency bands and associated behavioral and mental states are consistent with the advantages of synchronous logic systems. Cascaded oscillators can produce the frequencies found in EEGs.

##### 2.3.5.1. Synchronous logic systems

Logic systems have a timing problem in ensuring the various subcircuits change states in the correct chronological sequence. Synchronous logic systems generally have simpler circuit architecture and fewer errors than asynchronous systems. This is the reason nearly all electronic logic systems are synchronized by an enabling pulse to each component circuit. The enabling pulse in such systems is usually produced by an oscillator. The enabling input in Fig 3E and the oscillators in Fig 9 illustrate how such synchronization is possible with neural networks.

Timing problems are greater in sequential logic than in combinational logic, and greater in parallel processing than in serial processing. Much of the information processing in the brain involves sequential logic and nearly all of it is parallel. This means the selective pressure for synchronization in the brain would have been high, and the neural implementation proposed here is quite simple.

The processing speed in a synchronous system depends largely on the enabling oscillator’s speed. A large system like the brain that performs many diverse functions may have several different processing speed requirements. The trade-off for greater processing speed is a higher error rate. Functions that can tolerate a few errors, and that need fast results with many simultaneous small computations, require high processing speeds. Functions that are less dependent on speed or massive computation, or that require few errors, or whose component networks are large and complex and therefore slow to change state, call for slower processing.

##### 2.3.5.2. Synchronization and EEG frequency bands

The EEG frequency bands and associated behavioral and mental states are consistent with the function of multiple frequencies that was suggested in the preceding paragraph. Gamma waves (high frequencies) are associated with vision [53, 54] and hearing [14], which make sense out of massive data input in a few milliseconds. Beta waves are associated with purposeful mental effort [25], which may involve less data input while requiring few errors and complex operations. Alpha waves are associated with relaxed wakefulness [16], theta waves with working memory and drowsiness [25, 55], and delta waves with drowsiness and sleep [25]. These categories require successively slower information processing, and they have corresponding EEG bands of lower frequencies.

The high frequencies provided by the three-neuron ring oscillator and the wide variety of frequencies provided by cascaded toggles can produce this neural activity, as shown in Fig 11. The enabling signal from a neural oscillator (as illustrated in Fig 3E) can synchronize state changes in neural structures. The enabling pulse by itself does not produce state changes. When state changes do occur, the enabling pulse only ensures that they occur at regular times to avoid timing errors. So the initial ring oscillator’s high frequency signal could simply be connected directly and permanently to the enabling gates of networks in the visual and auditory cortexes, the first toggle’s signal connected to networks in the prefrontal cortex for purposeful mental effort, etc.

Providing synchronization for large numbers of neural structures performing many diverse brain functions would likely require more than a single cascade of oscillators. Only neural structures that are processing the same information need to be synchronized. Another set of structures processing different information could be synchronized by an oscillator in a different cascade. Because of the variation in delay times in different cascades’ initial ring oscillators, several cascades of oscillators could produce several different frequencies in each band simultaneously. A large number of neural structures synchronized in this way by many cascaded oscillators could exhibit the bands of matched periods found in EEGs.

## 3. Results and discussion

### 3.1. Explanations of known phenomena

#### 3.1.1. Short-term memory controversy

Cascaded oscillators and NFFs suggest a resolution to the question of whether short-term memory depends on neurons firing persistently or in brief, coordinated bursts [5, 6]: Memory is stored by persistent firing in flip-flops [4], and the coordinated bursts observed along with the persistent firing are due to the stored information being processed by several neural structures whose state changes are synchronized by a neural oscillator. An example of such short-term memory processing is a telephone number being reviewed in a phonological loop.

#### 3.1.2. Electroencephalography

##### 3.1.2.1. The cascaded oscillators hypothesis

The hypothesis that cascaded oscillators produce EEG frequencies is supported by the available data for neuron delay times and EEG frequency band peaks and boundaries, as shown in Fig 11. The oscillators’ designs and estimated frequency distributions graphed in Fig 11 can be derived directly from selective pressure for a biologically useful function without regard to EEG data. The useful function is synchronization of state changes in neural structures to avoid timing errors. In the tradeoff between speed and accuracy, the selective pressure is for a wide variety of enabling frequencies for diverse brain functions, including high frequencies for some information to be processed as fast as possible.

Cascaded oscillators provide a micro-level explanation of macro-level phenomena: The entire distribution of EEG frequencies in bands is determined by only two parameters - the mean and variance of neuron delay times.

EEG frequencies have two anomalous properties that are by-products of the cascaded oscillator solution to selective pressures, but otherwise have no apparent function: the octave relationship between EEG frequency bands and the close match between the distributions of EEG gamma frequencies and three-neuron ring oscillator frequencies. This makes it implausible that EEG phenomena are produced by a mechanism that is fundamentally different from cascaded oscillators.

##### 3.1.2.2. Answers to 16 questions raised by EEG phenomena

The cascaded oscillators hypothesis answers the questions in the section on unexplained EEG phenomena.

*What produces the widespread, synchronized, periodic firing?* 1) The firing is produced by cascaded oscillators. 2) The firing is periodic because neural structures are being enabled by oscillators. 3) The periodic firing is widespread because many neural structures are being enabled. 4) The firing is synchronized because a group of neural structures is being enabled by the same oscillator.

*What is the function of this widespread synchronization?* 5) The function of synchronization is timing error avoidance in processing information.

*What produces and what is the function of the wide distribution of EEG frequencies in bands?* 6) The frequencies occur in bands because each band is produced by a different oscillator. 7) The wide distribution of frequencies is due to the octave relationship between cascaded oscillators (100% exponential growth in periods with each successive oscillator by equation 5) and five oscillators. 8) The distribution of frequencies within each band is determined by the mean and variance of neuron delay times in the initial oscillators in the cascades (equations 6). 9) The function of the wide distribution of frequencies is meeting the needs of diverse brain functions in the trade-off between speed and accuracy.

*What produces the unimodal distribution in each band and the octave relationships between the peaks and boundaries?* 10) The unimodal distributions are due to the normal distribution of neuron delay times in the initial ring oscillators in cascades of oscillators. This makes the distribution of periods of each oscillator normal and the distributions of frequencies unimodal. 11) The ratio of consecutive boundaries and peak locations is 2 because consecutive cascaded oscillators increase the oscillation period by a factor of 2 (equations 5, 9, 10).

*What determines the specific frequencies of the peaks and boundaries?* The number of neurons in the ring oscillators must be the minimum of 3 to produce the high frequencies in the gamma band. 12) Equations 9 and 10 show the EEG band boundaries and 13) peaks are determined by this number (n = 3), the ring oscillators’ delay parameters μ_d_ and σ_d_, and the boundary or peak number i.

*Why do gamma oscillations peak at about 40 Hz?* 14) The three-neuron ring oscillator is the fastest neural ring oscillator. The estimated peak frequency from equation 10 is 38.4 Hz (illustrated in Fig 11).

*Why does the gamma band contain frequencies that are considerably faster than 40 Hz?*

The frequencies vary because of the variance in neuron delay times in the cascades’ initial oscillators. 15) As Fig 11 illustrates, all of the oscillator frequency distributions are skewed to the right, with the initial oscillator producing frequencies substantially greater than 40 Hz. In the particular estimate of Fig 11, 2% of the frequencies are greater than 75 Hz, and 0.4% are greater than 100 Hz.

*Why is there little agreement on the boundaries separating the EEG bands?* 16) The oscillators hypothesis implies that the estimates of EEG band boundaries are estimates of the intersections of the oscillators’ PDFs. This makes estimating boundaries difficult for two reasons.

The oscillators hypothesis implies that the probability of an EEG frequency being observed has a local minimum near each intersection of consecutive oscillator PDFs (Fig 11). This means that in a random sample of observed EEG frequencies, relatively few will be near the intersections. A small number of data points has a negative effect on the accuracy of estimates.

The overlapping oscillator PDFs (Fig 11) imply that the distributions of EEG frequencies associated with the various behavioral and mental states have overlapping ranges rather than discrete bands. Because two PDFs are equal at their intersection, a frequency near the intersection of two PDFs is almost equally likely to be produced by either of two oscillators. That is, an observed EEG frequency near a band “boundary” is almost equally likely to be observed along with the behavioral and mental state that defines the band on either side of the intersection. This makes obtaining accurate estimates of band “boundaries” especially difficult.

#### 3.1.3. A possible relationship between cascaded oscillators and epilepsy

Oscillations found in EEGs have a wide variety of frequencies, including high frequencies. These two properties necessitate a low tolerance for error in the duration of the input pulse to the oscillators proposed here. Irregularities such as variations in neuron delay times could cause serious errors in an oscillator’s output. Depending on the type of error that occurs, neural structures that are synchronized by the oscillator would either be disabled or enabled but unsynchronized. The resulting timing errors in neural firing and the brain’s efforts to deal with them may cause the abnormal electrical activity characteristic of epileptic seizures.

### 3.2. A simple, rigorous, statistical test of the cascaded oscillators hypothesis

#### 3.2.1. The data problem

Although the hypothesis that cascaded oscillators produce EEG phenomena is consistent with available data, as illustrated in Fig 11, the data are too imprecise for a rigorous statistical test of the hypothesis. The estimates found here for the neuron delay time parameters μ_d_ and σ_d_ were based on a description of the range of synapse delay times [25]. Available estimates of the EEG frequency bands’ peak frequencies are few and available only for three of the five major frequency bands. Estimates of band boundaries vary widely for reasons implied by the cascaded oscillators hypothesis as explained in the previous section. Estimates of both peaks and boundaries are routinely rounded to whole numbers. Some researchers do not even attempt to estimate a boundary separating two bands, instead giving a whole number frequency as the upper endpoint of one band and the next consecutive whole number as the lower endpoint of the next band. Estimates of means and variances of both neuron delay times and EEG frequency bands are apparently nonexistent.

#### 3.2.2. A simple test of the cascaded oscillators hypothesis from sampling data

A simple, rigorous test of the cascaded oscillators hypothesis is possible. All EEG phenomena predicted by the hypothesis follow from the main implication that the EEG bands and cascaded oscillators have the same distributions of frequencies. This implication can be tested statistically with random samples and the distribution relations of equations 6. As discussed previously, neuron delay times should be approximately normally distributed by the central limit theorem. This implies cascaded oscillator periods are also approximately normally distributed. A normal distribution is completely determined by its mean and variance. So it remains to be shown that EEG band periods are normally distributed and that the five main EEG band periods and five cascaded oscillator periods have equal means and variances.

The neuron delay time parameters μ_d_ and σ_d_ can be estimated from a random sample of neuron delay times. These estimates can be used to estimate the oscillator period distribution parameters μ_i_ and σ_i_ from equations 6. The mean and variance of the periods of one or more EEG bands can be estimated from a random sample of EEG periods (or frequencies). With standard tests for equal means and variances, the EEG estimates can be compared to the oscillator estimates of μ_i_ and σ_i_. The EEG sampling data can also be used to test EEG band periods for normal distributions. If the application of the central limit theorem to neuron delay times may be questionable, neuron delay times can also be tested for a normal distribution with the neuron delay time sampling data.

#### 3.2.3. Caveats

Because the oscillators’ frequency ranges overlap (Fig 11), the band to which an observed EEG period or frequency is assigned should be determined by the observed behavioral and mental state that defines a band, not by predetermined endpoints of bands. If EEG sampling data are measured in frequencies, they must be converted to periods before computing the sample mean and variance. (The period of the sample mean of frequencies is not the same as the sample mean of periods.) Sampling data should not be rounded to whole numbers. In using equations 6 to find the estimated oscillator parameters, recall that the value of n must be the minimum of 3. Sampling data for neuron delay times and EEG periods (or frequencies), or even estimates of means and variances, may already be available in some database.

Although it is possible that EEG frequencies are produced by cascaded oscillators with initial oscillators that are made up of specialized neurons whose delay times are different from the general population of neurons, this appears to be unlikely. Fig 11 shows the EEG frequency distributions are at least close to the values predicted by the general description of the range of neuron delay times that was used here to estimate oscillator neuron delay time parameters. Moreover, neurons in general and initial oscillator neurons in particular may have both evolved under selective pressure to function as fast as possible.

## 4. Acknowledgements

The simulations were done with MS Excel and CircuitLab. Network diagrams were created with CircuitLab and MS Paint. Graphs were created with MS Excel, Converge 10.0, and MS Paint. Predicted probabilities of high EEG frequencies were approximated with Converge 10.0. The author would like to thank Duncan Watson, Arturo Tozzi, David Garmire, Paul Higashi, Anna Yoder Higashi, Sheila Yoder, and especially Ernest Greene and David Burress for their support and many helpful comments.

## Notes

### Competing Interest Statement

The authors have declared no competing interest.

### Summary of Updates

The manuscript was substantially rewritten, with two major additions. 1. A possible connection to epilepsy was found. 2. The revision shows how a toggle flip-flop can maintain robust binary states in the presence of substantial levels of noise in the input pulse amplitude.

